# Newly Discovered migratory corridor and foraging ground for Atlantic green turtles, *Chelonia mydas*, nesting on Bioko Island, Equatorial Guinea

**DOI:** 10.1101/556191

**Authors:** Emily Mettler, Chelsea E. Clyde-Brockway, Shaya Honarvar, Frank V. Paladino

**Affiliations:** Department of Biology, Purdue University, Fort Wayne, Indiana, USA; Department of Biology, Purdue University, West Lafayette, Indiana, USA

**Author notes:** Current address: Department of Biology, Purdue University, Fort Wayne, Indiana, United States of America (Emily Mettler, Frank V. Paladino). These authors contributed equally to this work. These authors also contributed equally to this work (Chelsea E. Clyde-Brockway, Shaya Honarvar). Conceived and designed the experiments: EM, SH, FVP. Performed the experiments: EM, CECB. Analyzed the data: EM. Contributed reagents/materials/analysis tools: EM, SH, FVP. Wrote the paper: EM, CECB, SH, FVP.

## Abstract

This study is the first to use satellite telemetry to track post-nesting movements of endangered green turtles (*Chelonia mydas*) in the Gulf of Guinea. Satellite transmitters were attached in 2018 to six Atlantic green turtles nesting on Bioko Island, Equatorial Guinea, to track their post-nesting movements and locate their foraging grounds. Track lengths of 20-198 days were analyzed, for a total of 536 movement days for the six turtles. Migratory pathways and foraging grounds were identified by applying a switching state space model to locational data, which provides daily position estimates to identify shifts between migrating and foraging behavior. Turtles exhibited a combination of coastal and oceanic migrations pathways that ranged from 957 km to 1,131 km. Of the six turtles, five completed their migration and maintained residency at the same foraging ground near the coastal waters of Accra, Ghana until transmission was lost. These five resident turtles inhabit heavily fished and polluted waters and are vulnerable to a variety of anthropogenic threats. The identification of these foraging grounds highlights the importance of these coastal waters for the protection of the endangered Atlantic green turtle.

## Introduction

After entering the ocean as hatchlings, sea turtles spend the majority of their lives in the water, only emerging to lay eggs and or in some cases to bask [1] [2] [3]. Because of this, research on sea turtles has been largely restricted to nesting females, which has led to conservation efforts primarily focused on nesting beaches, rather than in-water habitat. Since in water habitats come with a variety of unique threats, including resource mining, fishing, and anthropogenic pollution, understanding oceanic habitat use and migration patterns is imperative to designing effective marine conservation strategies [4][5],[6],[7]. For example, endangered leatherback turtle populations in South Africa as well as Gabon have been unable to recover without in-water habitat protection, despite protection at nesting beaches, in part because of intensive long-line fishery operations off the coast of both countries with high rates of turtle bycatch [8],[9].

In West Africa, the Gulf of Guinea has experienced an increase in anthropogenic disturbances, putting the turtle population at risk from many threats including ship traffic, pollution, commercial and small-scale fishing operations [33], [5], and oil and gas development [7], [34]. Marine and coastal pollution in the waters of the Gulf of Guinea has caused a host of environmental threats, including oxygen depletion, faunal die-offs, and heavy metal and hydrocarbon accumulation in marine consumers [6], [35].

While sea turtles are protected under international and national law in most West African nations, incidental bycatch in fisheries operations is a major threat in the Gulf of Guinea. Green turtles are common bycatch in both gillnet and pelagic longline fishing operations [36], [37], [5]. Oil and gas development has also rapidly intensified in the Gulf of Guinea in recent years [34], and poses diverse, but difficult to measure, threats to sea turtle populations, with an increase in channel dredging, ship traffic, oil leaks, and chemical pollution, which can affect adult turtles that forage or travel close to offshore platforms [7]. These threats highlight the need to study migration patterns and foraging ground locations of sea turtles to better understand their vulnerabilities.

Adult green sea turtles have been known to migrate hundreds to thousands of kilometers between nesting seasons [38], [39], [40]. Generally post-nesting migrations are direct movements to foraging habitats, and turtles spend little energy on detours [41]; however, a number of studies have shown that in some cases individuals take indirect routes, including both open ocean and coastal pathways [42]. At foraging grounds green turtles generally maintain localized, near shore home ranges near sea grass beds [43], [44], [23]. Green turtles feed primarily on sea grass and other shallow water plant material; as such, there is limited food in the open ocean and direct migration to foraging grounds is thought to be advantageous by minimizing migration time and energy expenditure [45]. Just as green turtles show fidelity to nesting beaches [46], they also show fidelity to foraging grounds and post-nesting migratory routes are the same year after year [20]. Consequently, protecting migratory corridors and foraging grounds could have huge benefits for populations [40].

Little is known about the in-water movements and behavior of green turtles in the Gulf of Guinea. Green turtles that were flipper tagged on Bioko Island, Equatorial Guinea, in 1996-1998 have been recaptured in waters off the coast of Ghana, at least 1250 km from the nesting beaches of Bioko, in Corisco Bay, Gabon, about 280 km from Bioko, and off the coast of southern Gabon, at least 760 km from Bioko [47]. Since then, there have been no studies on post-nesting migration routes of green turtles from Bioko, and only one in the Gulf of Guinea, in which satellite telemetry was used to track green turtles nesting in Guinea-Bissau to their foraging ground off the coast of Mauritania [48].

To address the lack of knowledge on the post-nesting migratory routes of Atlantic green turtles in the Gulf of Guinea, we used satellite telemetry to track turtles from a nesting beach along the southern coast of Bioko Island, the second largest nesting rookery for green turtles within the Gulf [49], [50], [51]. Our specific objectives were to (1) map the post-nesting migration routes of green turtles from Bioko Island, (2) determine the directness of migratory routes and identify migratory corridors in the area, (3) categorize these migratory routes as coastal, open ocean, or both, and (4) locate coastal foraging grounds.

## Materials and methods

### Ethics Statement

This study was carried out in accordance with all federal, international, and institutional guidelines. All data was collected under the protocol approved by the Purdue Animal Care and Use Committee (PACUC Protocol Number 1410001142). Permissions to work within the protected area and with the study species were granted by the Instituto Nacional de Desarrollo Forestal y Gestión del Sistema de Áreas Protegidas (INDEFOR-AP permit #227), and the research protocol was approved by the Universidad Nacional de Guinea Ecuatorial (UNGE permit number 1011191091017).

### Study site

Bioko Island, Equatorial Guinea (2027 km^2^) is situated 175 km Northeast of mainland Equatorial Guinea. The southern coast has approximately 20 km of black sand beaches suitable for sea turtle nesting, all of which are within the legally protected Gran Caldera and Southern Highlands Scientific Reserve (Fig. 1). The remainder of Bioko’s 150 km coastline is generally unsuitable for sea turtle nesting due to cliffs, rocky beaches, and proximity to villages and roads [49]. Four species of sea turtles (leatherback, *Dermochelys coriacea*; green, *Chelonia mydas*; olive ridley, *Lepidochelys olivacea* and, hawksbill, *Eretmochelys imbricata*) nest across the five nesting beaches (8°66’-8°46’ E and 3°22’-3°27’ N), with the largest numbers of green turtle nests on beaches A, B, and C [51].This study was conducted on Beach C, chosen for its accessibility and high densities of green turtles (Fig. 1).

**Figure 1:**
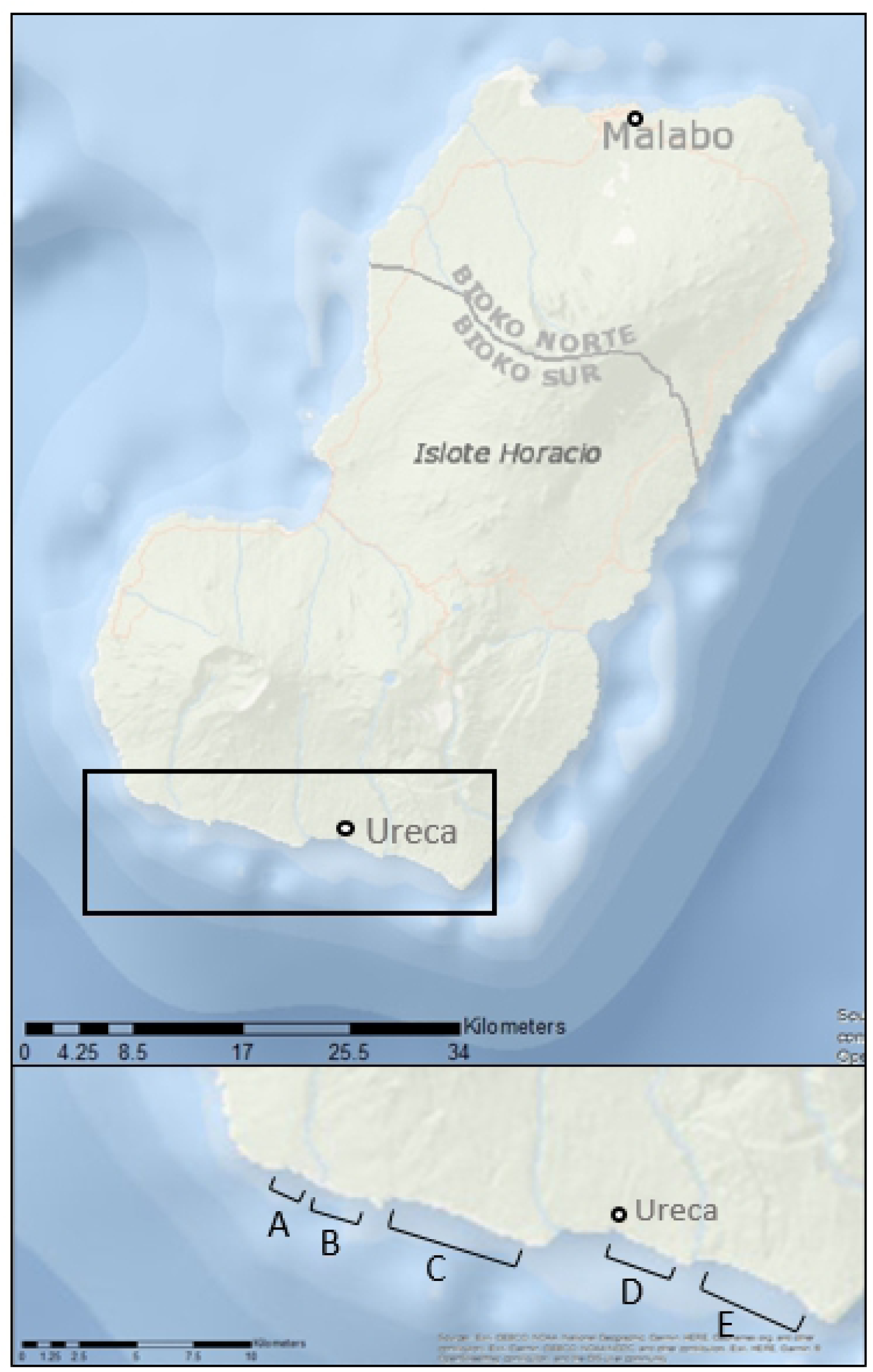
Map of the sea turtle nesting beaches on Bioko Island, Equatorial Guinea. Insert shows the five beaches (A-E) in relation to the nearest village, Ureca. Satellite transmitters were attached to green turtles nesting on Beach C, at the end of the nesting season in January-February 2018.

### Turtle Selection

Nesting season for green turtles on Bioko spans October through February [49]. Satellite transmitters were attached at the end of nesting season, in order to focus on tracking post-nesting migration and locational data from foraging grounds. Turtles that had laid their last nest, and therefore did not have developing vitellogenic follicles when scanned with a portable ultrasound (SonoSite 180 Plus; FUJIFILM SonoSite, Bothell, WA, USA), were preferentially selected as this generally indicates that the turtle is about to begin the post-nesting migration [13]. In addition, only turtles that had finished nesting and seemed to be in good health without any scarring or damage to the carapace where the transmitter would be attached were selected. Individuals were identified using a unique injectable passive integrated transponder (PIT) tag (AVID Identification Systems Inc., Norco, CA).

### Satellite transmitters

In January and February, 2018, six satellite transmitters (SirTrack, Kiwisat 202; Sirtrack, Havelock North, New Zealand) were attached to green turtles on Beach C, Bioko Island, after they had finished nesting. The transmitters were attached following the methods developed by Balazs et al. [52] modified by Luschi et al. [38], Troeng et al. [53], and Seminoff et al. [32]. Specifically, the carapace was cleaned, first with water, then with alcohol, and then scored with sandpaper to increase the strength of attachment. Transmitters were attached using Powers Pure50+ Two-Component Epoxy Adhesive (Powers, Brewster, NY, USA) to secure each transmitter to the second central scute of the carapace. Each turtle was restrained by a team of four or five researchers, and a wet cloth placed over the turtle’s eyes, to keep each turtle calm and in place while the epoxy hardened.

### Movement analysis

Location data was relayed via the Argos satellite system, and location points were filtered using the “argosfilter” package for R (R statistical software, R 3.4.3, Vienna, Austria), which removed any point that required a travel speed >5 km/hr [38]. The filtered location data was fit with a state-space model using the ‘bsam’ package [54] for R to estimate the behavioral state of the turtles. Filtered locational data was used instead of raw data to enhance the accuracy of the state space model [55]. The ‘bsam’ package, based on the Bayesian switching state space model developed by Jonsen et al. [56] was applied to the turtle tracks, using a hierarchical switching first-difference correlated random walk model (hDCRWS). The model was fit with a total of 5,000 Markov Chain Monte Carlo (MCMC) samples after 5,000 were discarded as burn-in, and every 10^th^ sample was retained. This model returns a behavioral mode of 1 (MCMC mean values <1.5) or 2 (values >1.5). Behavioral mode 1 is considered transiting behavior, and behavioral mode 2 is considered area restricted search (foraging) behavior.

Individual tracks were then mapped using ArcGIS 10.2 (Esri, Redlands, CA). Track length and daily travel distance were calculated using R from total track distance. Tracks were overlaid with ocean current data from the Ocean Surface Current Analysis Real-Time (OSCAR) from NASA [57].

## Results

All turtles (n=6) migrated west from the nesting beach. Tracks were analyzed for a total of 536 days. Average daily distance traveled was 49.5km, and the average total distance traveled for the five turtles that completed migrations was 1,055km. Two turtles exhibited oceanic migration routes and remained in transit across the Bight of Benin until reaching the coast of Togo and Ghana, where the state space model indicated a switch to foraging behavior (Fig. 2).

**Figure 2.**
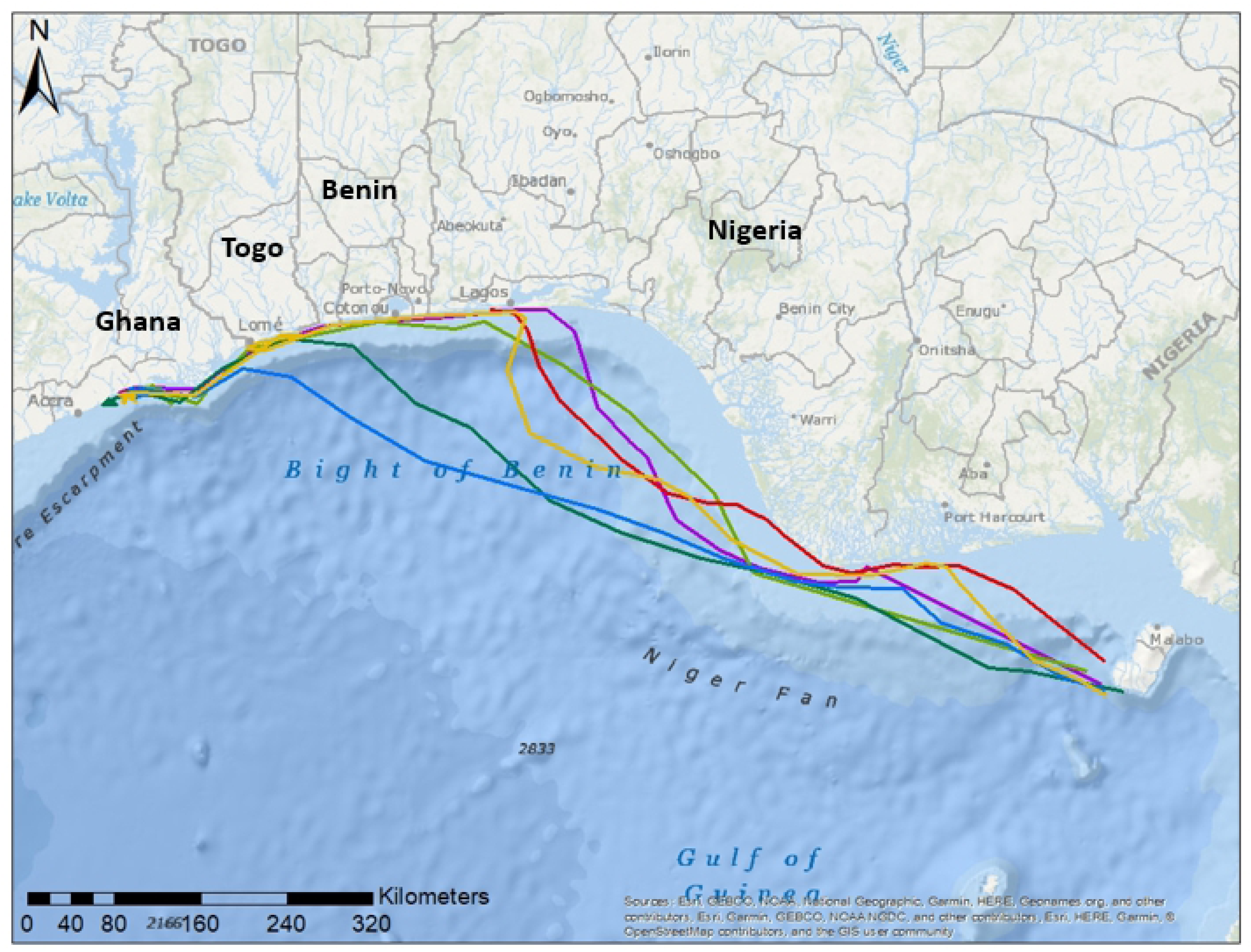
Post-nesting movements of six green turtles (Chelonia mydas) tracked from Bioko Island, after the 2017-18 nesting season. Individuals traveled an average of >1,000km using a combination of oceanic and coastal migratory routes. Two turtles exhibited oceanic migration routes (blue and dark green tracks); two turtles remained closer to the continental shelf (light green and purple tracks); two turtles migrated more directly across the Bight of Benin, to the coastal waters near Lagos, Nigeria, and then maintained a coastal route (yellow and red tracks).

These two turtles migrated for an average of 15 days and 989 km. The remaining four turtles used a combination of coastal and oceanic migratory routes, remaining closer to the continental shelf, a path that took an average of 23 days and 1098 km. Two of these turtles remained closer to the continental shelf, while two migrated more directly across the Bight of Benin, to the coastal waters near Lagos, Nigeria, and then maintained a coastal route (Fig. 2). Both oceanic and coastal migration routes remained in areas of weak currents for the duration of migrations (Fig. 3).

**Figure 3.**
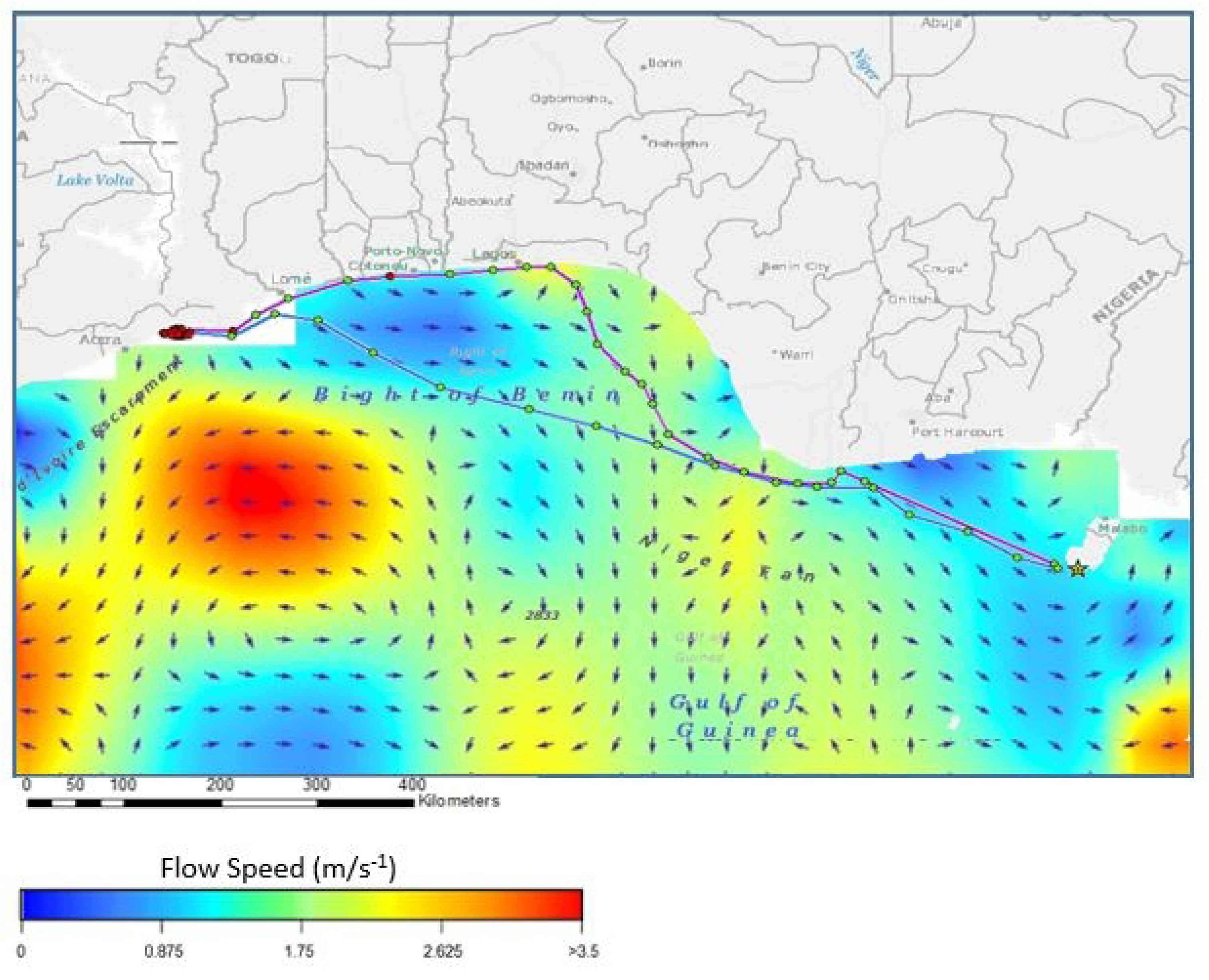
Ocean currents and daily locations (circles) of two green turtles tracked by satellite from Bioko Island across the Bight of Benin. A coastal and an oceanic migration route are overlaid onto ocean current data for the 5 day period from 15-20 Feb 2018. Green circles represent migrating behavior and red circles represent foraging behavior identified by the state-space model. Arrows represent current direction.

One turtle (Fig. 2: purple track) exhibited coastal migration and was in transit for 19 days until reaching the coastal waters of Lagos, Nigeria. Beginning on day 20, February 20^th^, all location transmissions were from land, in the Amuwo Odofin suburb of Lagos. Behavioral estimates provided by the state space model indicate that this turtle was still transiting, rather than foraging, when the transmissions from land began. As this turtle had no vitellogenic follicles remaining, there is no evidence that the turtle would have intentionally returned to land, and it is suspected that there was some human interaction that led to the transmitter being moved to land.

All five turtles ultimately began extended periods (>30 days) of residency and foraging behavior off the coast of Ghana, in a 50 km stretch east of Accra and west of the Volta River delta, after migration periods of 14-28 days (Fig. 4). Three turtles exhibited migrations interspersed with short (<3 days) periods of foraging off the coasts of Lagos, Nigeria, and Togo and Benin (Fig. 4). While the turtles exhibited both oceanic and coastal migrating behaviors, all exhibited near-shore foraging activity.

**Figure 4.**
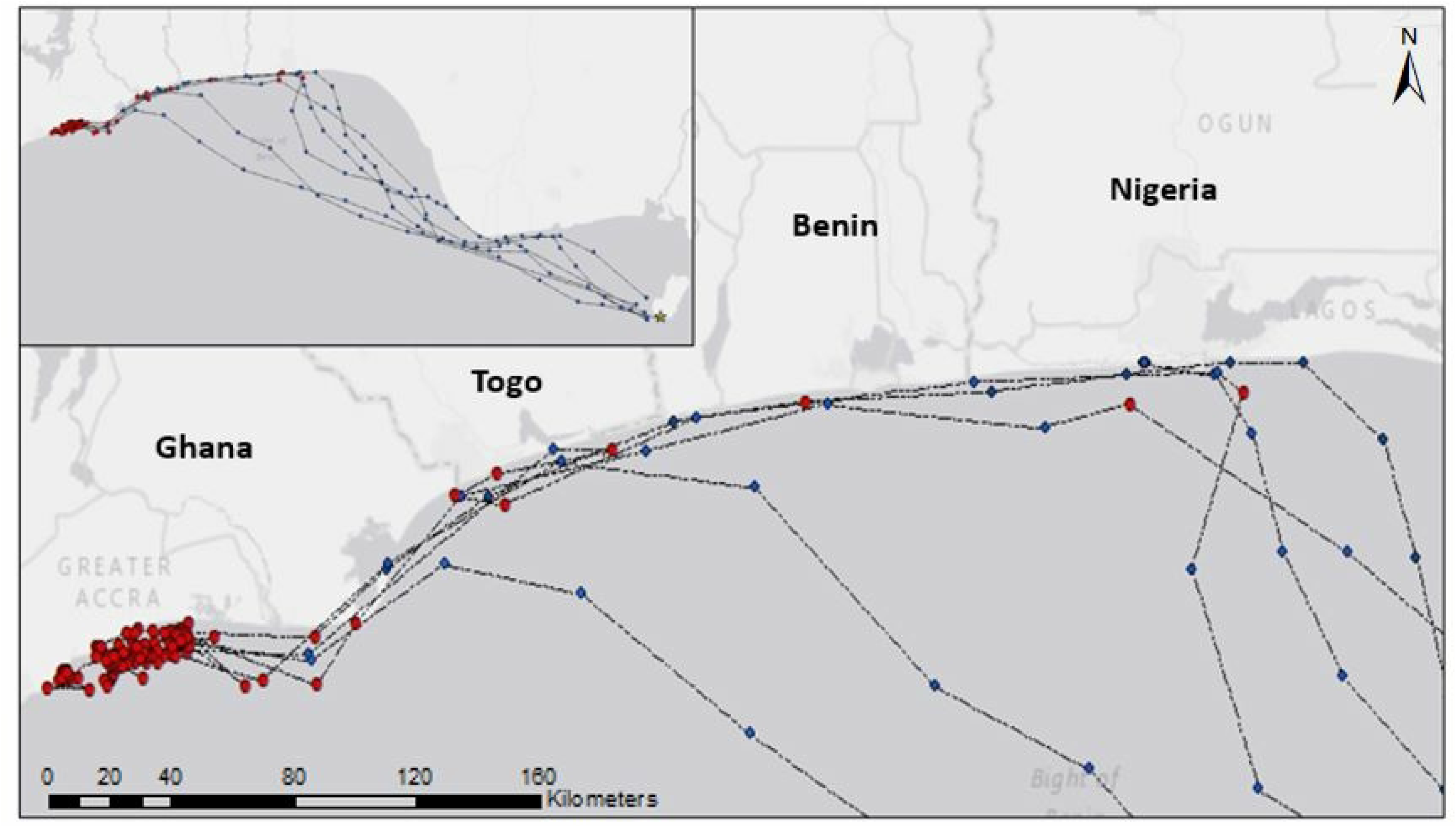
Daily locations (circles) of six turtles tracked from Bioko Island after the 2017-18 nesting season. Blue circles indicate transiting behavior and red circles indicate foraging behavior, as identified by the state space model. Three turtles exhibited migrations interspersed with short (<3 days) periods of foraging, while two exhibit direct migrations, followed by an extended period of foraging.

## Discussion

All six turtles migrated westward from Bioko Island, and five turtles completed their migration, ending at a previously undocumented foraging ground in the coastal waters of Ghana (Fig. 1). Previous recapture data suggested that some green turtles travel towards Ghana after nesting on Bioko, however the exact location of a foraging ground in the area was previously unknown [47].

Turtles exhibited both oceanic and coastal migration strategies, with two turtles traveling along a shorter route over deeper water (2000-3000m), and four traveling through shallower coastal waters (Fig. 1). Variations in migratory routes have been previously observed in green turtles nesting in Tortuguero, Costa Rica, as well as in the Galapagos [53], [32]. It has been hypothesized that these variations in migratory strategies may be linked to nutrient levels in post-nesting turtles, with more nutrient-depleted turtles taking the most direct route to a foraging ground, while turtle with higher nutrient levels may take a longer route [32].

Long distance migration is associated with high energy cost and all five complete migrations in this study were over 1,000 km. Turtles that used coastal migration routes exhibited short periods of foraging on the way to their final foraging ground (Fig. 2). Browsing behavior may decrease the overall energy cost of migration, and passing through suitable foraging habitat may be a benefit to a coastal migration pattern, mitigating the longer distance of coastal migratory routes. This behavioral plasticity has been documented in green turtles in the Caribbean and the coastal waters of Taiwan, in which turtles opportunistically utilized food sources that are available close to migratory routes [53], [42]. Similarly, turtles migrating from Bioko spent little time in areas that may have suitable foraging habitat, briefly foraging when advantageous and then continuing to a more distant foraging habitat, suggesting fidelity to a specific foraging ground. This could indicate that these stop over foraging habitats closer to Bioko cannot support sustained foraging of large numbers of turtles. It has also been reported that previously suitable coastal foraging habitat between Ghana and Nigeria has been compromised due to coastal erosion caused by sand mining and the development of harbors [33], which may necessitate longer migrations to more suitable habitat.

Green turtles often show fidelity to habitats, both nesting and foraging [46], [20]. Favoring specific foraging grounds, despite greater distances, may occur because of proximity to overwintering locations, resource limitations, territorial defense, or long-term fidelity to foraging grounds selected due to juvenile dispersal patterns [20]. However, foraging ground selection among these turtles cannot be explained by oceanic conditions, as both coastal and oceanic migration paths were against prevailing currents (Fig. 3). Given the low speeds of the currents, it is unlikely that countercurrent migrations greatly increased energy cost in this case.

Given the existence of nesting populations of green turtles on the beaches nearby this foraging ground in Ghana, and the apparent habitat suitability, it is likely that this foraging ground is used by multiple rookeries within the East Atlantic, including those nesting on Bioko Island [33], [58]. Furthermore, genetic studies have shown that green turtles that nest on Bioko Island share haplotypes with turtles found nesting in Ascension Island, Sao Tome, Principe, Corisco Bay, and Poilao [58]. Genetic similarities between populations suggests there are multiple populations using the same foraging and breeding grounds as those that nest on Bioko Island [58].

The discovery of this foraging ground is of particular importance, as only one other foraging ground used by green turtles in the Gulf of Guinea has been documented and protected-- Corisco Bay in Equatorial Guinea and Gabon. Yet all five turtles that completed migrations maintained residency in this newly discovered Ghanaian foraging habitat, highlighting the need for protection of this area. Migration routes passed through the exclusive economic zones (EEZs) of five countries, all of which rely heavily on fisheries for economic activity, which poses challenges to regulation and protection of this area. Only one of these countries has a marine protected area (MPA), and it lies well outside the migration corridor used by these turtles (Equatorial Guinea designated Corisco Bay as an MPA). Migrations passed through no MPAs, meaning throughout the migration pathways and within foraging grounds fishing is unrestricted. Studies have shown that green turtles in the Gulf of Guinea are caught as bycatch in both artisanal and industrial fisheries, in gillnets, driftnets, and purse and beach seines [59]. Although the extent of bycatch is difficult to quantify, it is suspected that mortality is significant, and is frequently underestimated by studies [60]. One of the six turtles involved in this study had a suspected interaction with humans after only 20 days of migrating, resulting in the transmitter being brought to land. While there is no way of knowing the nature of the interaction, turtles are consistently caught as bycatch in artisanal fishery operations in the area, and there is evidence that once caught, turtles are often transported to land and sold in markets [60].

Furthermore, this Ghanaian foraging ground lies near the outlet of a river that flows past the Kpone power plant as well as the Sakumo Lagoon, an important protected wetland heavily polluted by the inflow of industrial effluent, sewage, and domestic waste [61]. The Sakumo Lagoon has also been shown to have higher than average levels of Cadmium, Cobalt, Copper, Chromium, Nitrogen, and Zinc, which can have toxic effects on marine and aquatic wildlife [61].

## Conclusion

These threats highlight the need for further research into effects of fishing and pollution on this population, as well as the need to protect this valuable foraging habitat. Both industrial and domestic pollution as well as extensive commercial fishing are important issues when considering the protection of this newly discovered foraging ground. The distinct coastal foraging behavior of green turtles lends itself well to protection by spatially-explicit management strategies, such as zonal regulation of fishing and industrial dumping. Protecting nesting beaches in combination with delineating and protecting coastal foraging habitat on a national and multinational level may be key in conserving this highly migratory endangered species.

## Acknowledgments

The authors thank Lisa Sinclair for help with on the ground field operations, as well as research assistants with Purdue University Fort Wayne and the Bioko Marine Turtle Program, Brian Dennis, Amanda Rohr, Abby Khraling, and Sam Riley for help with transmitter attachment. We would also like to thank the National University of Equatorial Guinea (UNGE) and Instituto Nacional de Desarrollo Forestal y Gestión del Sistema de Áreas Protegidas (INDEFOR-AP) for their support. All attachment and animal handling procedures were made in compliance with IACUC protocol #1410001142. This study would not have been possible without financial support from Kosmos Trident Equatorial Guinea, Inc. grant. Funds from the Fort Wayne Children’s Zoo and the Sonoma County Community Foundation were also used to support this project.

## Supporting information captions

**S1 Dataset. Locational data from transmitter #107904**

**S2 Dataset. Locational data from transmitter #107905**

**S3 Dataset. Locational data from transmitter #107908**

**S4 Dataset. Locational data from transmitter #107910**

**S5 Dataset. Locational data from transmitter #107914**

**S6 Dataset. Locational data from transmitter #107915**

